# Decoding Attention Control and Selection in Visual Spatial Attention

**DOI:** 10.1101/2020.02.08.940213

**Authors:** Xiangfei Hong, Ke Bo, Sreenivasan Meyyapan, Shanbao Tong, Mingzhou Ding

## Abstract

Event-related potentials (ERPs) are used extensively to investigate the neural mechanisms of attention control and selection. The commonly applied univariate ERP approach, however, has left important questions inadequately answered. Here, we addressed two questions by applying multivariate pattern classification to multichannel ERPs in two spatial-cueing experiments (*N* = 56 in total): (1) impact of cueing strategies (instructional vs. probabilistic) and (2) neural and behavioral effects of individual differences. Following the cue onset, the decoding accuracy (cue left vs. cue right) began to rise above chance level earlier and remained higher in instructional cueing (∼80 ms) than in probabilistic cueing (∼160 ms), suggesting that unilateral attention focus leads to earlier and more distinct formation of the attentional set. A similar temporal sequence was also found for target-related processing (cued targets vs. uncued targets), suggesting earlier and stronger attention selection under instructional cueing. Across the two experiments, individuals with higher decoding accuracy during ∼460-660 ms post-cue showed higher magnitude of attentional modulation of target-evoked N1 amplitude, suggesting that better formation of anticipatory attentional state leads to better target processing. During target processing, individual difference in decoding accuracy was positively associated with behavioral performance (reaction time), suggesting that stronger selection of task-relevant information leads to better behavioral performance. Taken together, multichannel ERPs combined with machine learning decoding yields new insights into attention control and selection that are not possible with the univariate ERP approach, and along with the univariate ERP approach, provides a more comprehensive methodology to the study of visual spatial attention.

## 1. Introduction

Covert orienting of visual spatial attention can facilitate the processing of stimuli appearing at the attended location compared to that appearing at the unattended locations (Posner, 1980). The underlying neural mechanisms have been extensively studied by applying the event-related potential (ERP) technique to attention cueing paradigms (Dale, Simpson, Foxe, Luks, & Worden, 2008; Eimer, 2014; Eimer, van Velzen, & Driver, 2002; Grent-’t-Jong, Boehler, Kenemans, & Woldorff, 2011; Harter, Miller, Price, Lalonde, & Keyes, 1989; Hong, Sun, Bengson, Mangun, & Tong, 2015; Hopf & Mangun, 2000; Jongen, Smulders, & Van der Heiden, 2007; Kelly, Gomez-Ramirez, & Foxe, 2009; Lasaponara et al., 2018; Nobre, Sebestyen, & Miniussi, 2000; Yamaguchi, Tsuchiya, & Kobayashi, 1994). A series of cue-evoked slow ERP components have been identified that are higher over the hemisphere contralateral to the attended location, including early directing-attention negativity (EDAN), anterior directing-attention negativity (ADAN), late directing-attention positivity (LDAP) and biasing-related negativity (BRN). These ERP components respectively reflect early shift of attention, activation of attention control processes in frontal cortex and formation of facilitatory activity in target-specific visual cortex (Eimer, 2014; Grent-’t-Jong et al., 2011; Hopf & Mangun, 2000; Kelly et al., 2009; Lasaponara et al., 2018). Similarly, for target processing, the amplitude of several ERP components, including P1, N1, Nd1, Nd2 and late positive deflection (LPD), was enhanced for targets appearing at the cued location than that appearing at the uncued location, and this enhancement is thought to reflect the selection of task relevant stimuli for prioritized processing by attention (Curran, Hills, Patterson, & Strauss, 2001; Eimer, 1996; Eimer, 1998; Hong et al., 2015; Mangun & Buck, 1998; Mangun, Buonocore, Girelli, & Jha, 1998; Mangun & Hillyard, 1991; Rajagovindan & Ding, 2011; Talsma, Mulckhuyse, Slagter, & Theeuwes, 2007).

The univariate ERP approach, despite having generated a wealth of insights into the neural mechanisms of attention control and selection, has left some important questions inadequately addressed. For example, upon receiving the attention-directing cue, how long does it take to form the attentional control set? Do different cueing strategies (i.e., probabilistic vs. instructional) impact the timing of attention control? Predefined analysis windows used in previous univariate ERP research to measure the effect of attention vary significantly from study to study and do not yield precise answers to these questions. The impact of cueing strategies remains largely unexplored. In addition, there are significant individual differences in attention control and selection. Does stronger cue-triggered preparatory attention control lead to more effective selection of attended information? To what extent the attention selection of task-relevant stimuli is related to behavioral performance? Univariate ERP studies attempting to link cue-related ERPs with target-related ERPs (e.g., N1) and to link target-related ERPs with behavior (e.g., reaction time) have again yielded mixed results (Dale et al., 2008; Grent-’t-Jong et al., 2011; Harter et al., 1989; Talsma et al., 2007).

These issues may stem from the fact that univariate ERPs from single electrodes are not able to reflect the contributions of multiple neural processes taking place in distributed brain regions that collectively influence the subsequent neural or behavioral events in spatial attention (Eimer, 1998; Lasaponara et al., 2018; Mangun & Buck, 1998). For example, ADAN and LDAP both appear around 400-500 ms post-cue, but they each index different processes of preparatory attention and have distinct scalp topographies (Hong et al., 2015; Hopf & Mangun, 2000; Jongen et al., 2007; Lasaponara et al., 2018; Nobre et al., 2000). A multivariate approach taking into account the contribution of distributed neural processes that take place concurrently may provide a path forward to overcome the limitations of the univariate approach. Instead of treating different electrodes singly as in the univariate EEG/ERP approach, the multivariate pattern analysis/classification (also referred to as decoding) approach treats measurements from multiple ERP channels as a pattern representing the cognitive variable to be analyzed (Grootswagers, Wardle, & Carlson, 2017). To date, EEG-based decoding analysis has been applied to face detection (Cauchoix, Barragan-Jason, Serre, & Barbeau, 2014), working memory (Bae & Luck, 2018) and decision making (Bode et al., 2012). Here, we sought to apply this approach to understand the neural mechanisms of visual spatial attention control and selection. Our goal was to establish and compare the entire time courses of decoding for both cue-related (cue left vs. cue right) and target-related (cued target vs. uncued target) brain states across cueing strategies, and to link individual differences in decoding accuracy with attention enhancement of target processing and behavioral response.

Two cueing paradigms were considered in this study: probabilistic cueing (Mangun & Hillyard, 1991; Posner, 1980; Yamaguchi et al., 1994) and instructional cueing (Hopfinger, Buonocore, & Mangun, 2000; Snyder & Foxe, 2010; Worden, Foxe, Wang, & Simpson, 2000). The classic example of probabilistic cueing is the Posner paradigm in which the subject is encouraged to pay attention to a location where the target will be more likely to occur, but is required to respond to targets that are presented in both cued (valid trials) and uncued (invalid trials) locations (Posner, 1980). In this paradigm, there is strategic motivation to divide attention between both the cued location and the uncued location (Snyder & Foxe, 2010). In contrast, in instructional cueing, subjects are instructed to pay full attention to the cued location, and respond only to targets appearing at the cued location (ignoring targets presented in the uncued location). Compared with probabilistic cueing, this design produces a stronger attentional facilitation at the cued location, and has been adopted in many recent neuroimaging studies (Hong et al., 2015; Hopfinger et al., 2000; Liu, Bengson, Huang, Mangun, & Ding, 2016; Snyder & Foxe, 2010; Worden et al., 2000). Although both cueing strategies encouraged subjects to orient their attention to the cued location, because instructional cueing generates a more certain and a more focused unilateral attentional state than probabilistic cueing, we expect that decoding accuracy as a function of time during the cue-target interval and during the target processing would rise above chance level sooner and reach higher decoding accuracy for instructional cueing than for probabilistic cueing. We further predict that across individuals, the decoding accuracy during the cue-target interval, reflecting the strength of attention control, is positively associated with attention selection of the target and the decoding accuracy during target processing, reflecting the strength of attention selection, is positively associated with behavioral performance.

## 2. Material and Methods

### 2.1 Participants

The experimental protocols complied with the Declaration of Helsinki and were approved by the Institutional Review Board of Shanghai Mental Health Center (No. 2017-05R). Sixty-one healthy college students with normal or corrected-to-normal vision gave written informed consent and participated in this study. There were two experiments: Experiment 1 (N=32) utilized instructional cueing and Experiment 2 (N=29) utilized probabilistic cueing. Two participants were excluded in Experiment 1 due to poor data quality (i.e., less than 50% of trials remained after preprocessing). Three participants were excluded in Experiment 2 due to (1) poor task performance (N=1; accuracy < 50%) and (2) hardware issues (N =2; responses were not recorded correctly). Thirty participants were included in final analysis for Experiment 1 (10 females, aged between 18-25 years, all right-handed) and twenty-six participants were included in final analysis for Experiment 2 (10 females, aged between 20-24 years, all right-handed).

### 2.2 Stimuli and procedures

In both experiments, each trial began with an arrow cue (Experiment 1: 2.24° by 1.62°; Experiment 2: 2.29° by 1.62°) presented centrally for 200 ms, which pointed to either the left or right with equal probability. The cue was then replaced by a central fixation point (Experiment 1: a crosshair, 1.38° by 1.38°; Experiment 2: a dot, 0.57° by 0.57°) where subjects were required to maintain their fixation throughout each trial. Upon seeing the cue, subjects were instructed to shift their attention covertly to the cued direction. Two location markers (Experiment 1: squares, 2.39° by 2.39°, located 9.05° from the vertical meridian and 7.2° below the horizontal meridian; Experiment 2: dots, 0.57° by 0.57°, located 8.87° from the vertical meridian and 7.06° below the horizontal meridian) were presented in the left and right visual fields throughout the trials. After a cue-target interval of 1000-1200 ms, a target stimulus (Experiment 1: 1.67° by 1.67°; Experiment 2: 1.72° by 1.72°), either a plus sign or the letter ‘x’ with equal probability, was presented at one of the location markers. The target lasted for 200 ms and 100 ms in Experiment 1 and Experiment 2, respectively. In Experiment 1 (instructional cueing), the target was presented at the cued or uncued location with equal probability. Subjects were instructed to totally ignore the uncued location, and respond only to the plus sign appearing at the cued location. In Experiment 2 (probabilistic cueing), subjects were instructed to respond to the plus sign presented at both the cued (valid trials, 73.3%) and the uncued (invalid trials, 13.3%) locations. The remaining 13.3% trials in Experiment 2 were neutral trials with bilateral arrow cues, which did not provide any information about the location of forthcoming targets. The neutral trials were included for testing the behavioral effects of attention cueing and not included in the following ERP or decoding analysis. In both experiments, the inter-trial interval between the target offset and the cue onset of next trial was set at 2600 ms. Response to the plus sign was made by pressing a button on the response box with the right index finger as quickly and accurately as possible. Only responses made within 1600 ms after the target offset were considered as valid.

In Experiment 1, all stimuli were in black and presented in a white background (Fig. 1A). In Experiment 2, all stimuli were in white and presented in a black background (Fig. 1B). In both experiments, the paradigms were compiled and executed in the E-Prime 2.0 toolkit (Psychology Software Tools, Inc., Sharpsburg, USA), and all stimuli were presented on a 19-inch LCD monitor positioned 60 cm in front of the subject. Each trial block consists of 60 trials lasting for about 4-5 min. Subjects were first shown the experimental instructions, and then trained for at least 1 block to get familiarized with the task. After that, each subject finished 8 blocks, with a 2 to 3 min break between successive blocks.

**Figure 1.**
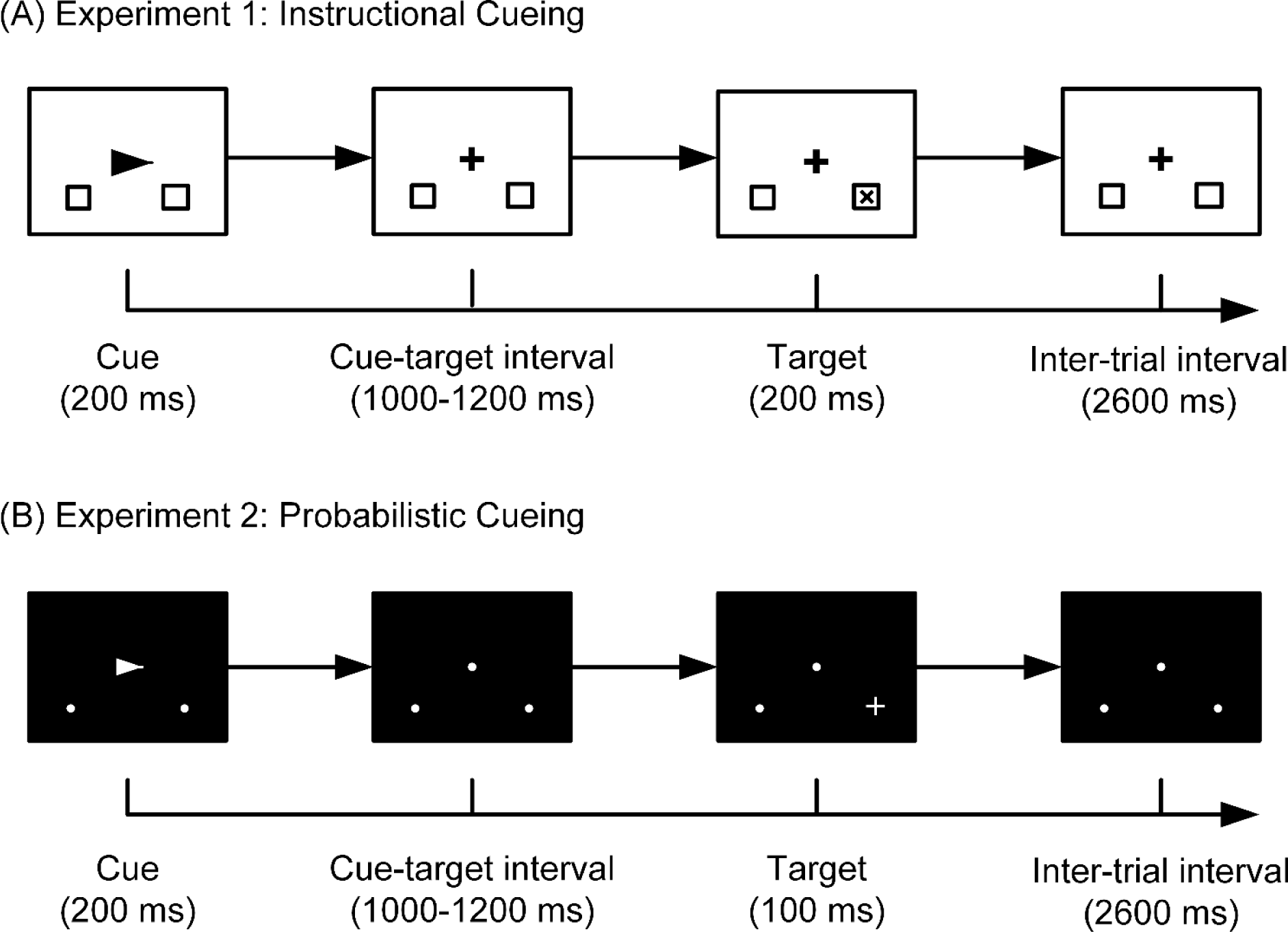
Experimental paradigms for the Instructional cueing dataset (*A*) and Probabilistic cueing dataset (*B*). In both paradigms, an arrow cue was first presented, directing the subject to covertly shift attention to either the left or the right lower visual field. Fixation was required throughout. After a random cue-target interval, a visual target (the letter ‘x’ or the plus sign) was presented in one of the location markers, and the subject responded to the target according to the paradigm requirements. In the instructional cueing paradigm, the subject was told to respond only to targets appearing at the cued location and totally ignore targets presented at the uncued location (50% probability). In the probabilistic cueing paradigm, the subject needed to respond to targets presented in both cued (73.3% probability, valid trials) and uncued (13.3%, invalid trials) locations. In the remaining 13.3% trials (neutral trials) in the probabilistic cueing paradigm, the cue was uninformative, consisting of a bilateral arrow. See Section 2.2 for more details.

### 2.3 EEG recording

EEG data were recorded using the BrainAmp MR Plus amplifier and EasyCap™ (Brain Products GmbH, Gilching, Germany). In Experiment 1, EEG signals were recorded from 32 electrodes (Fp1, Fp2, F3, F4, F7, F8, AFz, Fz, FCz, FC1, FC2, FC5, FC6, C3, C4, Cz, T7, T8, CP1, CP2, CP5, CP6, P3, P4, P7, P8, Pz, O1, O2, Oz, TP9, TP10), in which AFz and FCz were used as recording ground and reference, respectively. The sampling rate was set at 1000 Hz and the online anti-aliasing filter was set at 0.016-100 Hz. To monitor ocular activity, two additional electrodes were placed on the outer left ocular canthus and above the right eye recording horizontal and vertical electrooculograms (HEOG and VEOG). In Experiment 2, EEG signals were recorded from 65 electrodes (Fp1, Fp2, F3, F4, C3, C4, P3, P4, O1, O2, F7, F8, T7, T8, P7, P8, AFz, Fz, FCz, Cz, Pz, FC1, FC2, CP1, CP2, FC5, FC6, CP5, CP6, FT9, FT10, TP9, TP10, F1, F2, C1, C2, P1, P2, AF3, AF4, FC3, FC4, CP3, CP4, PO3, PO4, F5, F6, C5, C6, P5, P6, AF7, AF8, FT7, FT8, TP7, TP8, PO7, PO8, Fpz, CPz, POz, Oz), in which AFz and FCz were used as recording ground and reference, respectively. The sampling rate was set at 1000 Hz and the online anti-aliasing filter was set as 0.016-250 Hz. One additional electrode was placed below the right eye. In the offline analysis, we calculated the difference between this additional electrode below the right eye and Fp2 as the bipolar VEOG derivation, and the difference between FT9 and FT10 as the bipolar HEOG derivation.

### 2.4 EEG preprocessing

EEG preprocessing was conducted offline using EEGLAB and ERPLAB toolboxes, following the same general steps for both experiments. Specifically, continuous EEG data were first band-pass filtered into 0.1-40 Hz using a two-way Butterworth filter with zero phase shift (roll-off slope: 12 dB/oct). The power line noise was suppressed by a Parks McClellan notch filter at 50 Hz. Ocular artifacts were then corrected by independent component analysis (ICA) using the Infomax algorithm (Jung et al., 2000). Components related to eye movements are shown for three representative subjects in each experiment (Fig. 2). We then re-referenced EEG data to the average of the two mastoid electrodes (TP9, TP10). The original recording reference electrode was recalculated as FCz. After that, continuous EEG data were down-sampled to 250 Hz and then segmented into two types of epochs: one was time-locked to cue onset (from −1000 ms pre-cue to 1400 ms post-cue) and the other was time-locked to target onset (from −500 ms pre-target to 1000 ms post-target). It is worth noting that we extracted longer epochs than the periods we were interested in, so that we can minimize the filtering-related edge artifacts by trimming the two ends of the epoch (i.e., first and last 200 ms in both cue-related and target-related epochs; see below).

**Figure 2.**
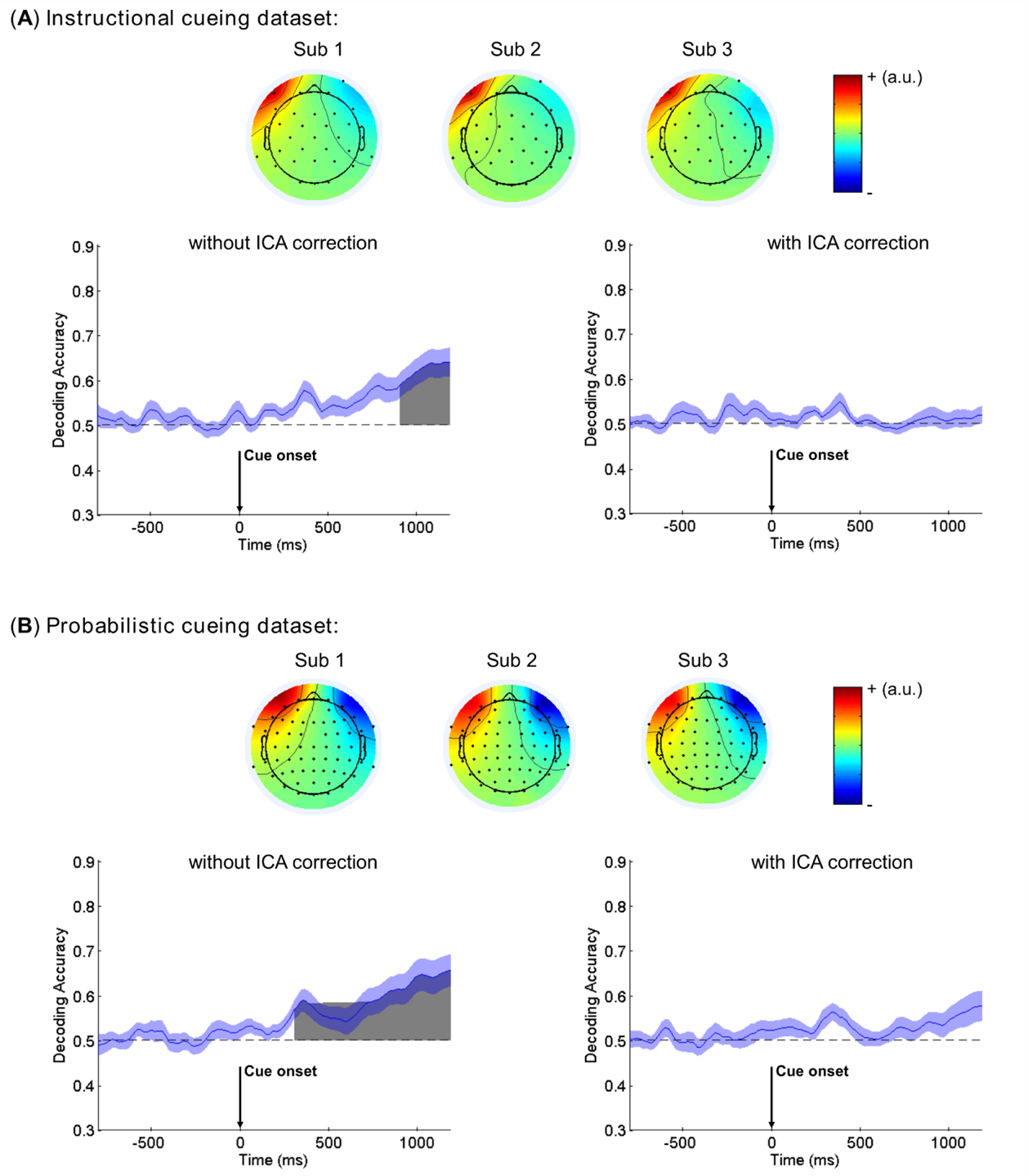
Removal of eye movement confounds. Independent components related to eye movements are shown for three representative subjects in each dataset. The decoding analysis was based on channels F7 and F8 which were near the eyes. Without ICA correction, the decoding is above chance-level during the late cue-target interval. Gray areas indicate clusters of time points in which the decoding was significantly greater than chance after the FDR correction for multiple comparisons. The blue shading indicates ±1 SEM. After removing eye movement related ICA component, the decoding accuracy returns to chance level.

Epochs meeting one or more of the following criteria were rejected: (1) the absolute value of maximal voltage difference over the whole epoch across all EEG channels exceeded 150 μV examined by a moving window (width: 200 ms; step: 50 ms) peak-to-peak function, (2) the absolute value of voltage on the whole epoch across all EEG channels exceeded 100 μV examined by a simple voltage threshold function, (3) epochs with any overt eye movements as detected by a moving window step function (width: 400 ms; step: 10 ms; threshold: 40 μV) based on HEOG amplitude, and (4) epochs with any overt eye blinks around the cue or target stimuli presentation period (−200-200 ms) as detected by a moving window peak-to-peak function (width: 200 ms; step: 10 ms; threshold: 50 μV) based on VEOG amplitude.

The trial rejection rates across subjects (mean ± standard deviation) of cue-related epochs were 17.73% ± 14.00% (cue left) and 17.48% ± 14.02% (cue right) for Instructional cueing dataset, and 19.75% ± 14.69% (cue left) and 19.86% ± 15.50% (cue right) for Probabilistic cueing dataset. There were no significant differences in trial rejection rates between cue left and cue right in either dataset (both *p* > 0.7, paired sample *t*-test). The trial rejection rates of target-related epochs were 7.27% ± 7.59% (attended targets) and 8.13% ± 7.88% (ignored targets) for Instructional cueing dataset, and 11.89% ± 12.92% (valid targets) and 11.64% ± 13.32% (invalid targets) for Probabilistic cueing dataset. There were no significant differences in trial rejection rates between cued (attended or valid) and uncued (ignored or invalid) targets in either dataset (both *p* > 0.4, paired sample *t*-test). Only epochs with correct behavioral performance that were also artifact-free after correction in all channels were included in the following analysis.

### 2.5 Univariate ERP analysis

Conventional univariate ERP analysis was performed by averaging EEG epochs of the same condition triggered either by the cue or by the target. Specifically, cue-related epochs were averaged according to the cue direction (left, right) with pre-cue interval (−200, 0 ms) as baseline, yielding cue-related ERPs for each condition, electrode and participant. Target-related epochs were averaged according to the target location (left, right) and attention (cued, uncued) with pre-target interval (−200, 0 ms) as baseline, yielding target-related ERPs for each condition, electrode and participant.

### 2.6 Multivariate pattern classification/decoding

#### 2.6.1 Overview

We examined whether multichannel patterns of ERPs can be used to reveal neural representation of visual spatial attention. Since single-trial EEG data are noisy and attention-related ERPs have small amplitudes (e.g., typically less than 1 μV for EDAN, ADAN and LDAP), it would be difficult to decode their patterns based on single-trial EEG data. However, averaging across trials can substantially improve the signal-to-noise ratio, and thus increase decodability (Grootswagers et al., 2017; Steven J Luck, 2014). Thus, we applied a recently proposed ERP-based decoding approach (Bae & Luck, 2018, 2019), in which multiple EEG epochs from a given cue or target condition were first averaged to yield the ERPs at each channel, and decoding was then performed on multichannel patterns of ERPs rather than on single-trial data, as is often the case in EEG-based decoding studies. Decoding accuracy was computed based on the average of a test set of epochs with the same condition that was not in the training set used to define the classifier.

In addition to ERPs that are phase-locked to the event of interest (cue onset or target onset), EEG data also contain non-phase-locked activity related to visual spatial attention, such as alpha oscillations (Hong et al., 2015; Liu et al., 2016; Worden et al., 2000). However, alpha oscillation was not the focus of this study. To ensure that our decoding was driven by ERP activity rather than alpha oscillations, we applied low-pass filtering at 8 Hz using the EEGLAB eegfilt() routine before decoding. The first and last 200 ms in both cue-related and target-related epochs were removed to minimize edge artifacts due to filtering.

To increase the efficiency of decoding analysis and reduce computation time, we further down-sampled the data to 50 Hz (1 data point every 20 ms). For cue-related epochs, we obtained a 4-dimensional data matrix for each participant, with dimensions being time (100 time points), cue direction (left vs. right), trial, and electrode site (29 electrodes for Instructional cueing dataset, 60 electrodes for Probabilistic cueing dataset). For target-related epochs, we also obtained a 4-dimensional data matrix for each participant, with dimensions being time (55 time points), target condition (cued vs. uncued), trial, and electrode site.

#### 2.6.2 Support vector machine (SVM) classifier

The classifier was based on linear SVM and trained through the MATLAB fitcsvm() function. The decoding procedure at a given time point included a training phase and a testing phase. Training and testing phases were based on different trials. Specifically, a three-fold cross-validation procedure was applied at each time point. The data from 2/3 of the trials (randomly selected) were used to train a classifier (training), and then the performance of the classifier was assessed with the data from the remaining 1/3 of trials (testing). For the cue period, we first organized cue-related epochs with respect to the cue direction (left vs. right) and then divided all trials of the same condition into three equal-sized groups. One or two trials from each cue direction was omitted if the trial number is not evenly divisible by 3. The trials in each group were averaged to yield a scalp distribution of ERPs for the time point being analyzed (a matrix of 3 groups × 2 cue directions × 29/60 electrodes). We then performed z-score normalization across channels at each time point to eliminate possible baseline differences between conditions. This normalization was performed within each group to prevent any data leakage. The data from two of the three groups served as a training dataset, which were used to train a SVM classifier, and the data from the remaining group served as a testing dataset. The trained SVM classifier was then used, with the help of the MATLAB function predict(), to predict the direction of visual spatial attention for the testing dataset. The output of this function provided a predicted cue direction for each observation in the testing dataset. Decoding accuracy was then computed by comparing the true labels of cue direction with the predicted labels. Decoding was considered correct only if the classifier correctly determined the direction of cued attention (left or right). The chance performance was 50%.

This decoding procedure was repeated three times, once with each of the three groups of data serving as the testing dataset. The entire procedure as described above was iterated 20 times, each time with a new random assignment of trials into three groups. This iteration could help to minimize idiosyncrasies associated with trial assignments, and thus yield a more stable result. After that, decoding accuracy was collapsed across 2 cue directions, 3 cross-validations, and 20 iterations, yielding an averaged decoding accuracy for a given time point based on 120 decoding attempts (2 cue directions × 3 cross validations × 20 iterations). After this procedure was applied to each time point from −800 to +1200 ms (relative to cue onset), the averaged decoding accuracy values were smoothed across time points to minimize noise using a 5-point moving window (equivalent to a time window of ±40 ms).

For the target period, we organized target-related epochs with respect to the condition of attention deployed to the target (cued vs. uncued), collapsed across left and right visual hemifields. SVM was again used to classify the attention condition based on the spatial distribution of target-related ERP signals over the scalp. The decoding procedure was identical to that for the cue period, except that the time period of analysis was the 55 time points from −300 to +800 ms (relative to target onset). Decoding was considered correct only if the classifier correctly determined the condition of attention deployed to the target (cued or uncued). The chance performance was 50%.

In addition to decoding accuracy, we also examined the extent to which different channels drove the classifier performance by reconstructing the spatial distribution of the transformed classifier weights, namely, the activation patterns or weight maps. This was obtained by multiplying the classifier weights with the covariance matrix of the original data, yielding the weight maps for the classifier at each time point (Haufe et al., 2014).

#### 2.6.3 Statistical analysis of decoding accuracy

If the multivariate ERP patterns across electrodes contain information about the two conditions being compared (cue left vs. cue right for cue-related activity; cued target vs. uncued target for target-related activity), then the decoding accuracy should be greater than chance level, which was 50%. We tested whether the group-level decoding accuracy at each time point was above chance level by performing one-tailed signed rank test against the chance level of 50%. False Discovery Rate (FDR) was used for correcting the multiple comparison problem with *q* < 0.05. Furthermore, to control for possible isolated time points that might have high decoding accuracy by chance, we removed time windows with less than 3 contiguous time points that survived FDR correction.

#### 2.6.4 Minimizing the impact of eye movements

Although trials with overt eye movements were rejected and ICA-based artifact correction was applied to remove voltage fluctuations generated by micro eye movements, to ensure that the decoding results were not contaminated by eye movements, we conducted an additional set of decoding analyses based on two EEG channels near the eyes (F7/F8), and compared the results between ICA-corrected data and uncorrected data. As suggested by Quax, Dijkstra, van Staveren, Bosch, and van Gerven (2019), if the decoding accuracy contained contributions from eye movements, we should observe above chance level decoding during the cue-target interval even using the ICA-corrected data.

As expected, if we did not perform ICA correction, the decoding accuracy was above chancel level during the cue-target interval in both datasets (Fig. 2), indicating that systematic eye movements contributed to the decoding even after excluding trials contaminated by overt eye movements. After ICA correction, however, the decoding accuracy remained at chance level throughout the cue-target interval in both datasets, suggesting that the ICA correction for eye movements was successful and the decoding analysis reported below was not adversely impacted by eye movements.

### 2.7 Correlating decoding accuracy with attention modulation of target processing and behavior

#### 2.7.1 Cue-related period

The target-related N1 component (∼170 ms after stimulus onset) is one of the most extensively reported electrophysiological measures of visual spatial attention (Hillyard & Anllo-Vento, 1998; Hong et al., 2015; S. J. Luck, Woodman, & Vogel, 2000; Mangun & Hillyard, 1991). In this study, we calculated the differences of N1 amplitudes between cued targets and uncued targets (with the negative sign maintained) as the index of attention modulation, and correlated it with the decoding accuracy in the cue-target interval using Pearson correlation (2-tailed). This correlation analysis was performed at each time point from 0 to +1200 ms (relative to cue onset) to yield a time course of *r* values and corresponding *p* values.

Although the paradigms of the two experiments differed in terms of whether or not targets presented in the uncued location required response, subjects were expected to voluntarily shift their attention to the cued location in both paradigms. In other words, the general attention orienting process during the cue-target interval is the same between the two datasets, and the cue-target interval is also of the same duration in the two datasets (1000-1200 ms). When appropriate, we performed a meta-analysis by combining the *p* values from the two datasets using the Liptak-Stouffer meta-analysis, a well-validated method for combining multiple datasets (Huang & Ding, 2016; Liptak, 1958; Liu et al., 2017). Specifically, the *p* value of correlation at each time point was converted to its corresponding *Z*-score using the equation: *Z*_i_ = *Φ*^−1^(1 − *p*_i_), where *Φ* is the standard normal cumulative distribution function and *i* represent the *i*^th^ dataset. Here *i* = 1 (Instructional cueing dataset) or 2 (Probabilistic cueing dataset). Next, a combined *Z*-score for the correlation at each time point was calculated using the Liptak–Stouffer formula:

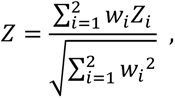

where *Z* is the meta-analysis *Z*-score across the two datasets, *Z*_i_ is the *Z*-score of the *i*^th^ dataset, *W*_i_ = √N_i_ is the weight of the *i*^th^ dataset, and *N* is the number of subjects in the *i*^th^ dataset. Finally, the combined *p* value was identified from the meta-analysis *Z*-score at each time point.

Next, we assessed the statistical significance of the correlation results from the above procedure by performing a cluster-based permutation test. Specifically, we first found clusters of contiguous time points for which the combined single-point correlation was significant (meta-analysis *p* < 0.05) and computed the cluster size (numbers of contiguous time points). The minimal cluster size was set as 1. We then asked whether a given cluster size was greater than the size that would be expected by chance through permutation tests. This controls the Type I error rate at the cluster level, yielding a probability of 0.05 that one or more clusters would be significant if true decoding accuracy was at chance (Groppe, Urbach, & Kutas, 2011). To determine whether a cluster size was larger than expected by chance, we generated a null distribution of cluster size values via permutation tests. The permutation was performed at the stage of between-subject correlation rather than at the stage of within-subject SVM training or testing. Specifically, after obtaining the mean decoding accuracy and N1 modulation of each subject, we shuffled the indices of subjects before performing the Pearson correlation in each iteration of the permutation. This was designed to generate the correlation results that would be obtained by chance if the decoding accuracy was not related to N1 modulation. It should be noted that we applied the same shuffled indices of subjects for all the time points in a given iteration, instead of using different shuffled indices for each time point independently. This was implemented to preserve the temporal auto-correlation of the continuous EEG data (Bae & Luck, 2019; Linkenkaer-Hansen, Nikouline, Palva, & Ilmoniemi, 2001). In each iteration, the above permutation was performed in each dataset separately, and the *p* values at each time point from the two datasets were then combined using the Liptak-Stouffer meta-analysis introduced above. After that, we computed the cluster size for which the combined single-point correlation was significant (*p* < 0.05) based on the meta-analysis *p* values for that permutation iteration. If we observed more than one cluster with significant combined *p* values, we then took the largest cluster size as the size for that iteration.

The above procedure was iterated 1000 times to produce a null distribution for the cluster size. To compute the *p* value for a given cluster size observed in the actual datasets, we simply found where this *p* value fell within the null distribution of cluster size. The *p* value for a given cluster was then set based on the nearest percentiles of the null distribution. If the obtained cluster size is larger than the maximum of permuted cluster size, we then reported *p* < 0.001. If an observed cluster size was in the top 95% of the null distribution, we rejected the null hypothesis and concluded that the correlation was significant for that observed cluster.

Finally, to see whether univariate ERPs during the cue-target interval were able to predict the attention modulation effect of target processing, we performed the same correlation analysis between univariate ERPs and N1 attention modulation, and used the same permutation test to assess the statistical significance. This was performed on the basis of ERP difference waves (cue left *minus* cue right) for each channel separately. FDR correction was applied to correct for multiple comparison across channels. The corrected *p* values were then used for find clusters of contiguous time points. Since the channel numbers differed between the two datasets, we reported the results (the cluster-level *p* value and mean *r* value in each cluster) for each dataset separately instead of combining the results using meta-analysis.

#### 2.7.2 Target-related period

To reveal the functional significance of ERP-based decoding during the target-related period, we correlated the mean decoding accuracy with behavioral performance. For the Instruction cueing dataset, due to the lack of behavioral cueing metrics, we used the mean RT to attended targets as the behavioral measure. For the Probabilistic cueing dataset, we calculated the differences of RT between valid and invalid (invalid *minus* valid) trials as the behavioral measure. Because the behavioral metrics were different between the two datasets, we reported the results for each dataset separately instead of combining them through meta-analysis.

The correlation analysis and permutation test for statistical significance were the same as that used in the cue-related period, except that we used different time windows for the correlation analysis. Specifically, this correlation analysis was performed at each time point from 0 to +560 ms (relative to target onset) in the two datasets separately, yielding an *r* value and a *p* value at each time point in each dataset. The reason we chose this time period was because the average RT was around 470 ms in the Instructional cueing dataset, and the average RT to valid targets was around 500 ms in the Probabilistic cueing dataset. Thus, 0-560 ms provided adequate coverage of the neural processes leading from target processing to behavioral response.

Finally, like the cue-related period, we performed the same correlation analysis between univariate ERPs and behavioral performance, and used the same permutation test to assess the statistical significance. This was performed on the basis of ERP difference waves (cued target *minus* uncued target) for each channel separately. The FDR-corrected *p* values were used for find clusters of contiguous time points, the same as correlation analysis for cue-related ERPs. Because the channel numbers and behavioral metrics used differed across the two datasets, we reported the results (the cluster-level *p* value and mean *r* value in each cluster) for each dataset separately instead of combining the results using meta-analysis.

## 3. Results

### 3.1 Behavioral results

#### Instructional cueing dataset

Accuracy and reaction time (RT) were analyzed separately for the cue left and cue right trials. Accuracy was defined as the percentage of correctly performed trials, and RT was averaged across all correctly performed trials that required responses. Paired sample *t*-test suggested no differences in accuracy (cue left: 99.55% ± 0.08% vs. cue right: 99.49% ± 0.11%; *t*_(29)_ = 0.504, *p* = 0.618) or RT (left targets: 467.85 ± 9.91 ms vs. right targets: 471.38 ± 10.01 ms; *t*_(29)_ = −1.038, *p* = 0.308) between cue left and cue right trials. These results demonstrated that the subjects deployed visual spatial attention according to the paradigm design.

#### Probabilistic cueing dataset

Accuracy and RT were calculated in the same way as in the Instructional cueing dataset. Paired sample *t*-test suggested no differences in accuracy between different types of trials (valid: 98.04% ± 0.67%, invalid: 98.54 % ± 0.51%, neutral: 97.89% ± 0.72%; all *p* > 0.05). The analysis of RT (valid: 501.76 ± 17.88 ms, invalid: 566.81 ± 22.04 ms, neutral: 512.13 ± 18.14 ms) suggested that subjects responded more slowly in invalid trials than in valid trials (valid vs. invalid: *t*_(25)_ = −5.587, *p* < 0.001). Furthermore, the RT in neutral trials was longer than that in valid trials (neutral vs. valid: *t*_(25)_ = 2.426, *p* = 0.023), and shorter than that in invalid trials (neutral vs. invalid: *t*_(25)_ = −5.437, *p* < 0.001). These results, consistent with previous reports, demonstrated that the subjects deployed visual spatial attention according to the paradigm design.

### 3.2 Attentional modulation of target-related N1

#### Instructional cueing dataset

We conducted a two-way ANOVA with Attention (attend vs. ignore) and Target Location (left vs. right) as within-subject factors on target-related N1 amplitudes. We observed the main effect of Attention (*F*_(1,29)_ = 60.114, *p* < 0.001), suggesting that attention significantly modulated the sensory processing of targets (see Fig. 3A). No main effect or interaction was observed for the factor of Target Location.

**Figure 3.**
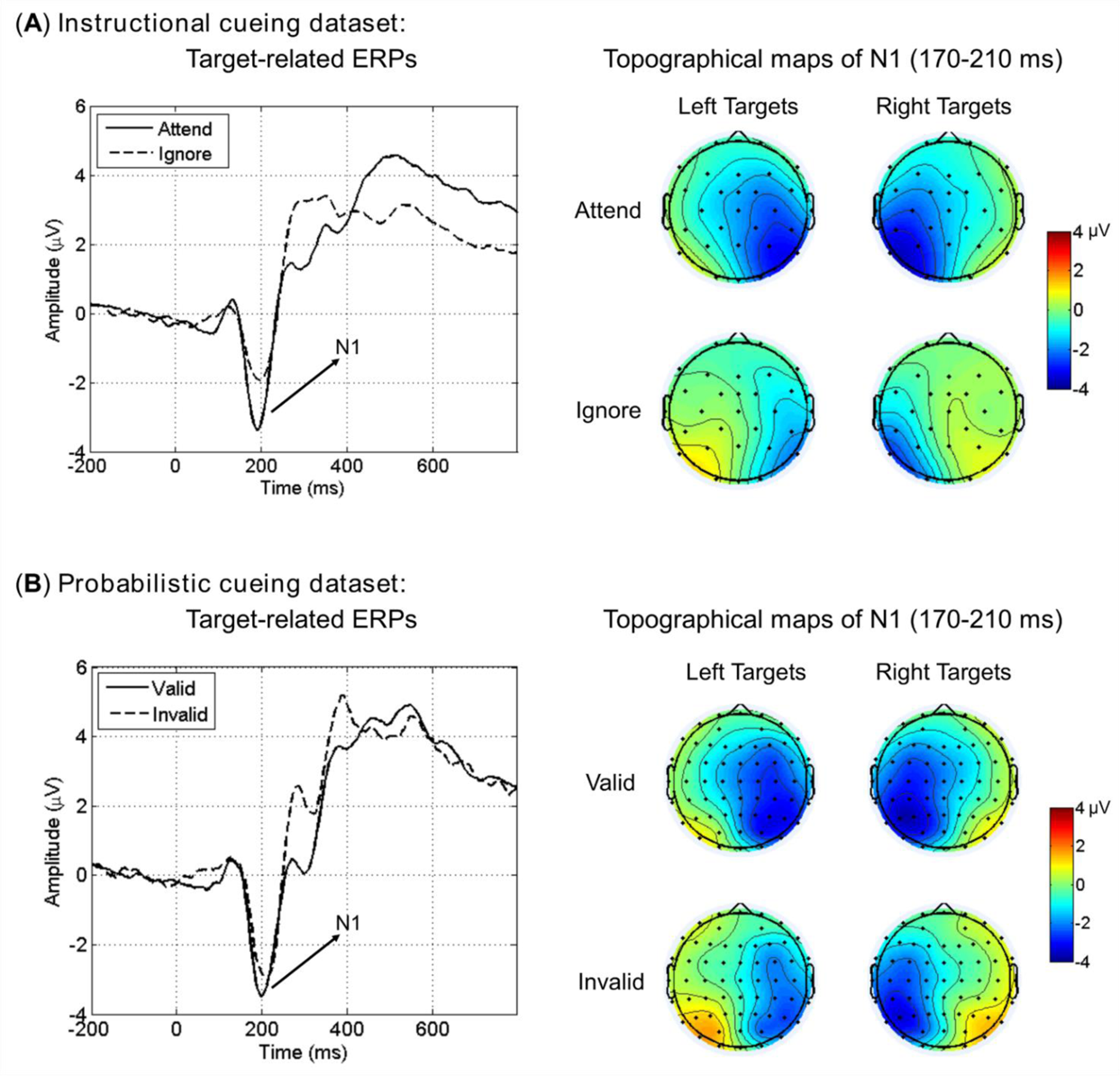
Target-related ERPs and attentional modulation on N1 amplitudes in the Instructional cueing dataset (*A*) and Probabilistic cueing dataset (*B*). Target-related ERPs were constructed over posterior scalp regions that were contralateral to target location (i.e., left hemisphere for right targets, right hemisphere for left targets), then averaged across left and right hemispheres.

#### Probabilistic cueing dataset

A similar two-way ANOVA was carried out with Attention (valid vs. invalid) and Target Location (left vs. right) as within-subject factors on target-related N1 amplitudes. Again, we observed the main effect of Attention (*F*_(1,25)_ = 8.418, *p* = 0.008), suggesting a significant attentional modulation of the sensory processing of targets (see Fig. 3B). No main effect or interaction was observed for the factor of Target Location.

#### Comparing N1 modulation between the two datasets

We compared the magnitude of attentional modulation of N1 amplitudes (cued target *minus* uncued target) between the two datasets using *t*-test. Results showed that N1 modulation was stronger in Instructional cueing dataset than in Probabilistic cueing dataset (−1.23 ± 0.16 μV vs. −0.65 ± 0.22 μV, *t*_(54)_ = −2.159, *p* = 0.035).

### 3.3 Decoding attention control in the cue-related activity

#### 3.3.1 Decoding accuracy

##### Instructional cueing dataset

As Fig. 4A illustrates, the decoding accuracy began to rise above chance level (50%) shortly after the cue onset, and more specifically, a signed rank test (FDR-corrected) indicated that the decoding was significantly greater than chance level starting at ∼80 ms and remaining significant until the end of the cue-target analysis interval. The weight maps showed a frontally-posteriorly lateralized distribution during the 400-600 ms interval, which then diminished as time progressed, and became a centrally lateralized distribution in the later cue-target interval (1000-1200 ms). These weight maps were similar to the ERP topographical maps in corresponding time windows (See Figure S1 in Supplemental Information).

**Figure 4.**
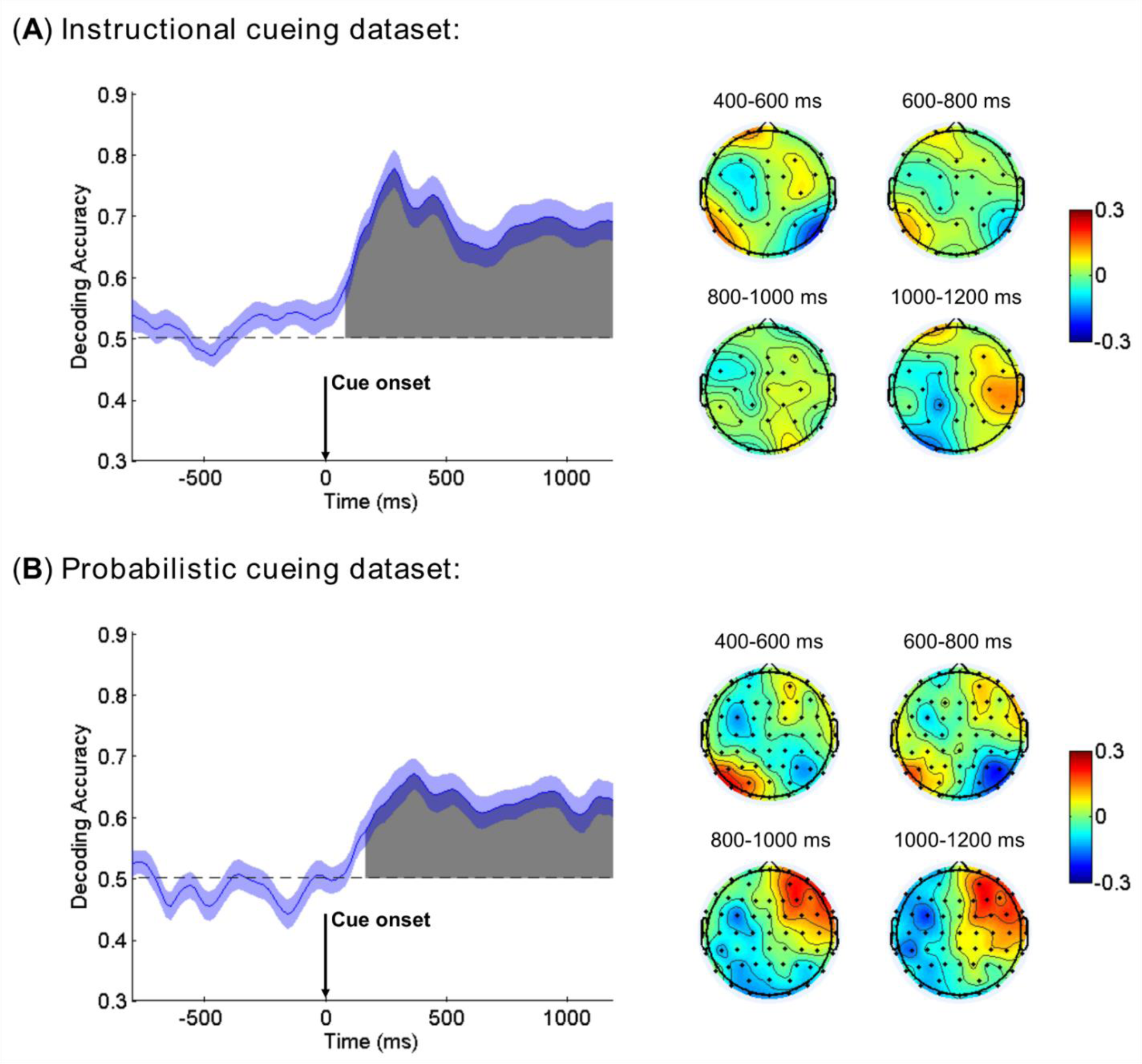
Mean accuracy of ERP-based multivariate decoding for cue-related neural processing (cue left vs. cue right) in the Instructional cueing dataset (*A*) and Probabilistic cueing dataset (*B*). Chance level performance (0.5) is indicated by the horizontal dash lines. Gray areas indicate clusters of time points in which the decoding was significantly greater than chance after the FDR correction for multiple comparisons. The blue shading indicates ±1 SEM. Weight maps from successive time points within the indicated windows were averaged and shown on the right.

##### Probabilistic cueing dataset

As Fig. 4B illustrates, the decoding accuracy began to rise above chance level (50%) at ∼160 ms after the cue onset according to a signed rank test (FDR-corrected), and remained significant until the end of the cue-target analysis interval. Overall, the weight maps from probabilistic cueing were consistent with that from instructional cueing, except that in the later cue-target interval (1000-1200 ms), the weight maps from probabilistic cueing were more anteriorly distributed.

##### Comparing the onset of above chance decoding between the two datasets

We used bootstrap resampling (100 times) across subjects to compute a distribution of decoding onset times for each dataset. A *t*-test suggested that the decoding onset time was significantly earlier in Instructional cueing dataset than in Probabilistic cueing dataset (64.8 ± 3.4 ms vs. 161.4 ± 3.5 ms, *t*_(198)_ = −19.706, *p* < 0.001).

##### Comparing the decoding accuracy between the two datasets

We divided the cue-target interval after decoding onset into two equal-sized time windows, i.e., 200-700 ms (early) and 700-1200 ms (late), and averaged the decoding accuracy within each window for each subject and dataset. A two-way ANOVA with Dataset (Instructional vs. Probabilistic) as a between-subject factor and Window (early vs. late) as a within-subject factor revealed a main effect of Dataset (*F*_(1,54)_ = 4.649, *p* = 0.036), suggesting significantly higher decoding accuracy for Instructional cueing dataset than for Probabilistic cueing dataset. No other main effect or interaction was observed for this ANOVA.

#### 3.3.2 Linking decoding accuracy and attentional modulation of target processing

As with any other neurophysiological variables, decoding accuracy varies significantly across individuals. We take that as an opportunity to examine the functional significance of decoding accuracy. For cue-related decoding accuracy, we correlated it with the magnitude of the attentional modulation of the target-evoked N1 component, the classic marker of attention selection. The correlation coefficients (*r* values) and corresponding *p* values for Instructional cueing dataset and Probabilistic cueing dataset are shown in Fig. 5A and Fig. 5B as functions of time. We observed negative correlations (i.e., higher decoding accuracy predicts larger N1 attentional modulation) during the cue-target interval for both datasets, and such correlation reached statistical significance around ∼500 ms; this finding was consistent across the two datasets. For Probabilistic cueing dataset, the direction of this correlation was reversed briefly around ∼900 ms, but the same was not observed in Instructional cueing dataset.

**Figure 5.**
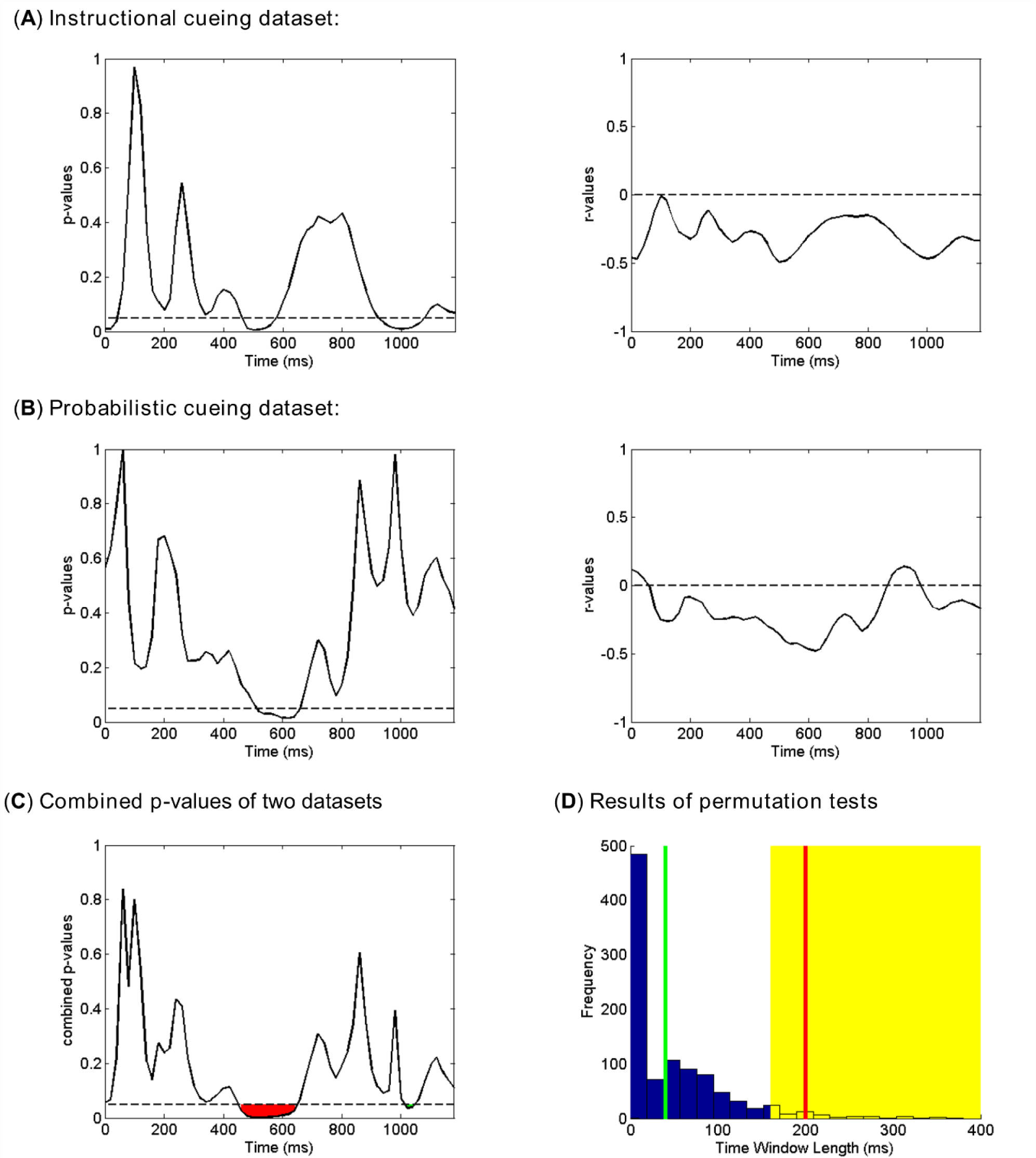
Results of between-subject correlation between cue-related decoding accuracy and attentional modulation of target-related N1 in Instructional cueing dataset (*A*) and Probabilistic cueing dataset (*B*). The *p* values from the two datasets were combined by the Liptak-Stouffer meta-analysis (*C*). Panel *D* shows the results of permutation tests for the two consecutive time windows identified in Panel *C*. The null distribution was estimated from 1000 permutations of the data, by randomly pairing one subject’s decoding accuracy with another subject’s N1 modulation. If the window length from the observed data (red and green lines) falls within the top 5% of values from the null distribution (indicated by the yellow area), the observed window is considered to be significant.

The correlation results from the two datasets were further combined through the Liptak-Stouffer meta-analysis. The correlation between decoding accuracy within the 460-660 ms post-cue window (see Fig. 5C, red region) and N1 attention modulation was significantly (*p* < 0.05) negative, i.e., higher decoding accuracy predicted greater N1 attention modulation. Further permutation test confirmed that the correlation during this ∼200 ms length window was statistically significant (*p* = 0.037) (see Fig. 5D, red line). There was another shorter window (1020-1060 ms post-cue) showing significant correlation (see Fig. 5C, green region). However, permutation test suggested that the correlation during this ∼40 ms length window was not significant (*p* = 0.443) (see Fig. 5D, green line).

Next, we examined whether cue-related univariate ERPs predicted the N1 attention effect. The univariate ERP difference waves (cue left *minus* cue right; see Figure S1 in Supplemental Information) was correlated with the attentional modulation of N1, and the results were shown (0-1200 ms post-cue) in Fig. 6. FDR correction was first applied to correct for multiple comparison across channels, and permutation test was then applied to find significant clusters of contiguous time points. No significant cluster was identified for either dataset. Thus, unlike the decoding accuracy derived from multichannel ERPs, individual differences in univariate ERPs showed no relation to that in attentional modulation of N1.

**Figure 6.**
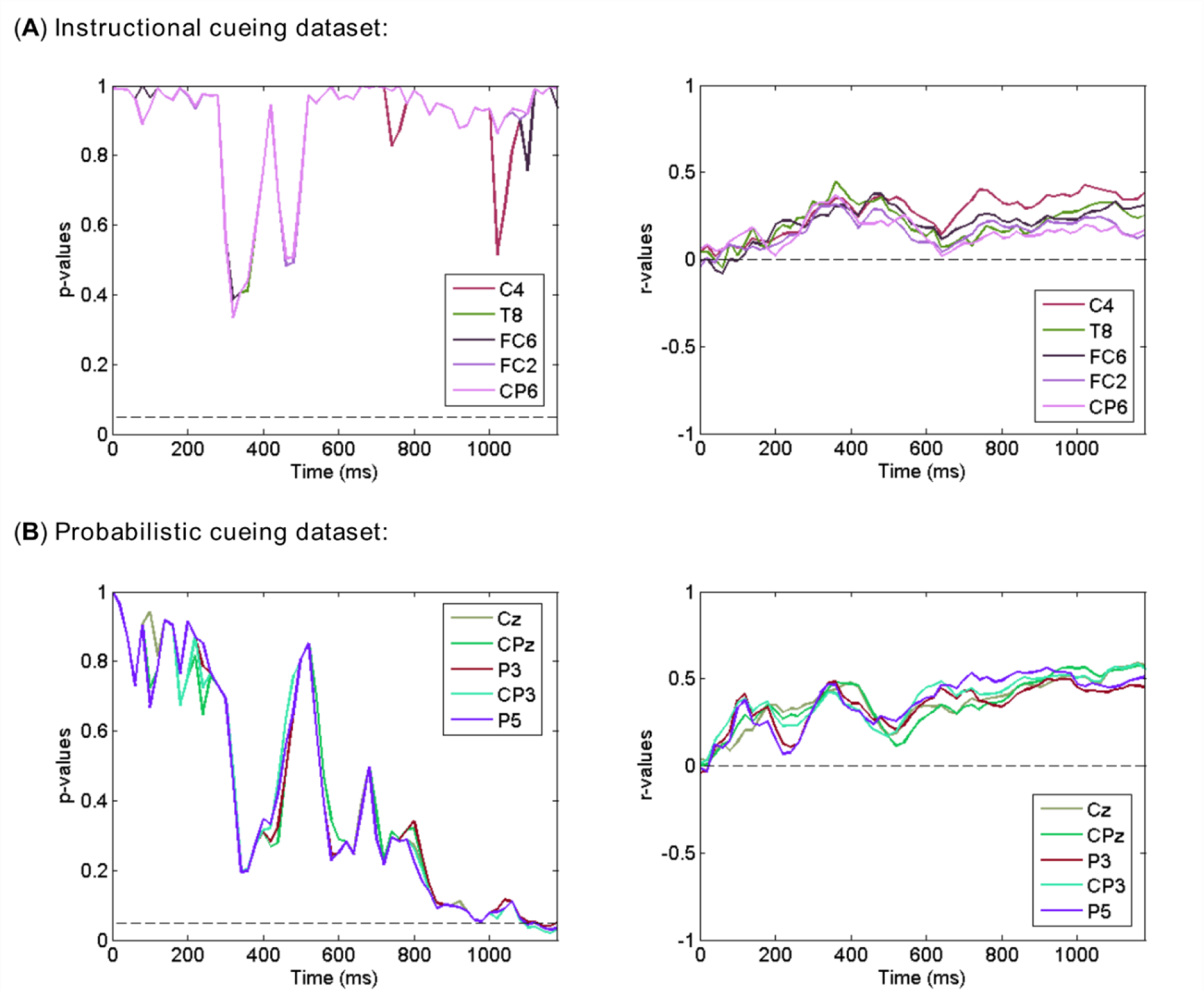
Results of between-subject correlation between cue-related univariate ERP difference wave (cue left *minus* cue right) and attentional modulation of target-related N1 in Instructional cueing dataset (*A*) and Probabilistic cueing dataset (*B*). 5 channels with the smallest averaged *p* values within 0-1200 ms interval were shown. All *p* values were FDR-corrected for channels at each time point. No significant effects were found.

### 3.4 Decoding attention selection in the target-related processing

#### 3.4.1 Decoding accuracy

##### Instructional cueing dataset

After the target onset, as Fig. 7A illustrates, the decoding accuracy began to rise above chance level (50%) at ∼100 ms, and remained significant above chance level for the remainder of the analysis period (mean RT was ∼470 ms). The weight maps showed a frontal distribution during the 200-400 ms interval, and became more parietal-distributed as time progressed. These weight maps were similar to the ERP topographical maps from corresponding time windows (See Figure S2 in Supplemental Information).

**Figure 7.**
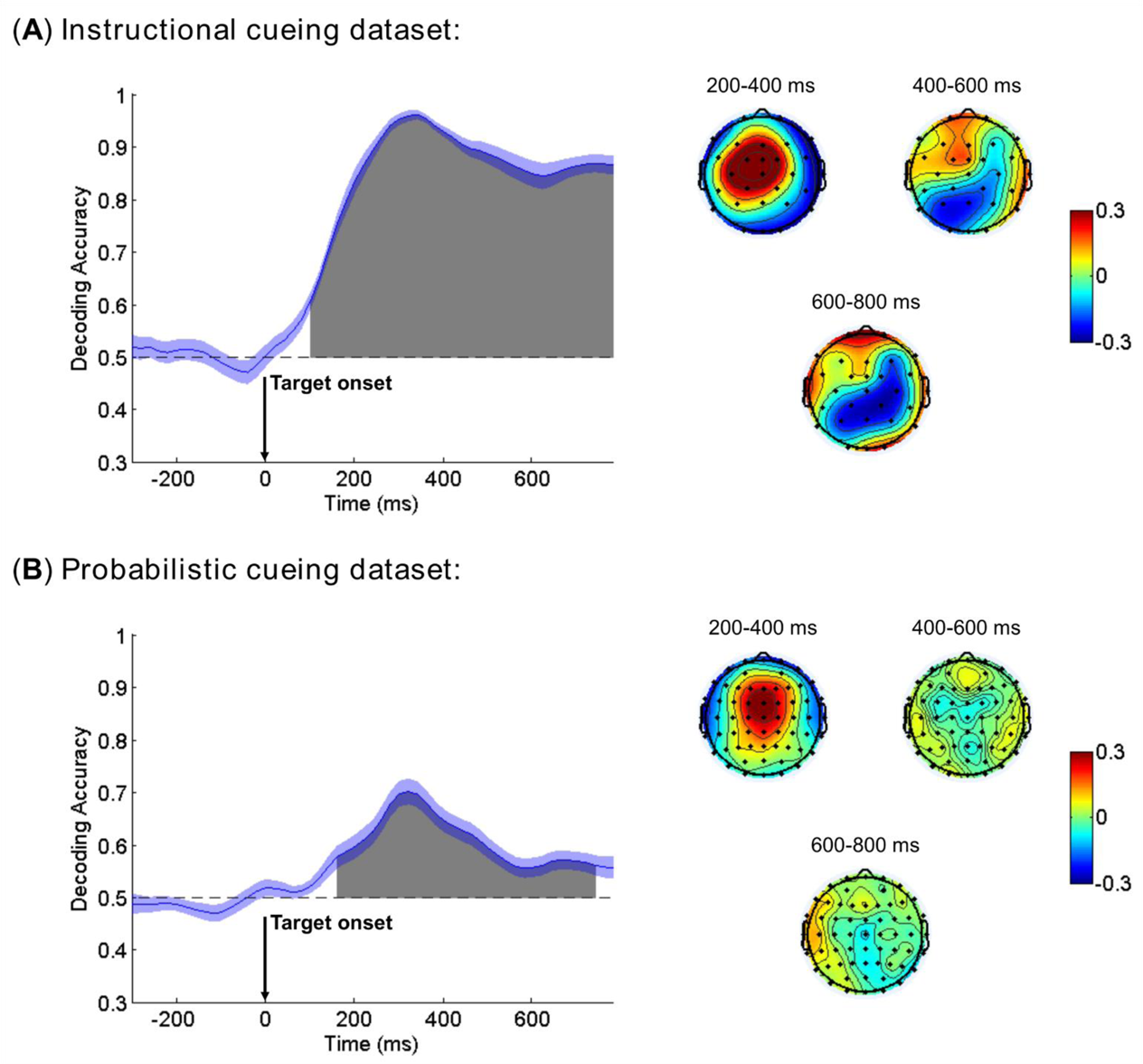
Mean accuracy of ERP-based multivariate decoding for target-related epochs (cued vs. uncued) in Instructional cueing dataset (*A*) and Probabilistic cueing dataset (*B*). Chance level performance (0.5) is indicated by the horizontal dash lines. Gray areas indicate clusters of time points in which the decoding was significantly greater than chance level after the FDR correction for multiple comparison problem. The blue shading indicates ±1 SEM. Weight maps from successive time points within the indicated windows were averaged and shown on the right.

##### Probabilistic cueing dataset

Similarly, as Fig. 7B illustrates, the decoding accuracy began to rise above chance level (50%) at ∼160 ms after the target onset. The decoding accuracy gradually decreased after reaching its peak at ∼300 ms, but still remained significant until 740 ms (mean RT in invalid trials was ∼566 ms). Similar to the ERP topographical maps (See Figure S2 in Supplemental Information), the weight maps showed a frontal distribution during the 200-400 ms interval, which then diminished as time progressed.

##### Comparing the onset of above chance decoding between the two datasets

We used bootstrap resampling (100 times) across subjects to compute a distribution of decoding onset times for each dataset. A *t*-test suggested that the decoding onset time was significantly earlier in Instructional cueing dataset than in Probabilistic cueing dataset (85.8 ± 3.0 ms vs. 156.0 ± 4.1 ms, *t*_(198)_ = −13.734, *p* < 0.001).

##### Comparing decoding accuracy between the two datasets

We divided the target analysis interval after decoding onset into two equal-sized time windows, i.e., 200-500 ms (early) and 500-800 ms (late), and averaged the decoding accuracy within each window for each subject and dataset. A two-way ANOVA with Dataset (Instructional vs. Probabilistic) as a between-subject factor and Window (early vs. late) as a within-subject factor revealed a main effect of Dataset (*F*_(1,54)_ = 187.032, *p* < 0.001), suggesting significantly higher decoding accuracy for Instructional cueing dataset than for Probabilistic cueing dataset. A main effect of Window (*F*_(1,54)_ = 39.033, *p* < 0.001) suggested that decoding accuracy significantly declined in the late window than in the early window. No interaction was observed.

#### 3.4.2 Linking decoding accuracy with reaction time

To examine the functional significance of decoding accuracy following the target onset, we correlated individual differences in decoding accuracy at each time point within 0-560 ms post-target interval with individual differences in behavioral performance (RT to attended targets for Instructional cueing dataset, RT difference between invalid and valid trials for Probabilistic cueing dataset). The correlation coefficients (*r* values) and corresponding *p* values for Instructional cueing dataset and Probabilistic cueing dataset are shown as functions of time in Fig. 8A and Fig. 8B. From 180 to 560 ms, the correlation between decoding accuracy and RT was significant (*p* < 0.05) for Instructional cueing dataset. Permutation test confirmed that the correlation during this window was statistically significant (*p* < 0.001) (see Fig. 8A, red line), and the negative *r* values suggested that individuals with higher decoding accuracy had faster responses to attended targets. For Probabilistic cueing dataset, the correlation between decoding accuracy and RT difference was significant (*p* < 0.05) from 60 to 340 ms. Permutation test confirmed that the correlation during this window was statistically significant (*p* = 0.004; see Fig. 8B, red line), and the positive *r* values suggested that individuals with higher decoding accuracy exhibited larger RT differences between invalid and valid trials (i.e., stronger benefits from attention cueing).

**Figure 8.**
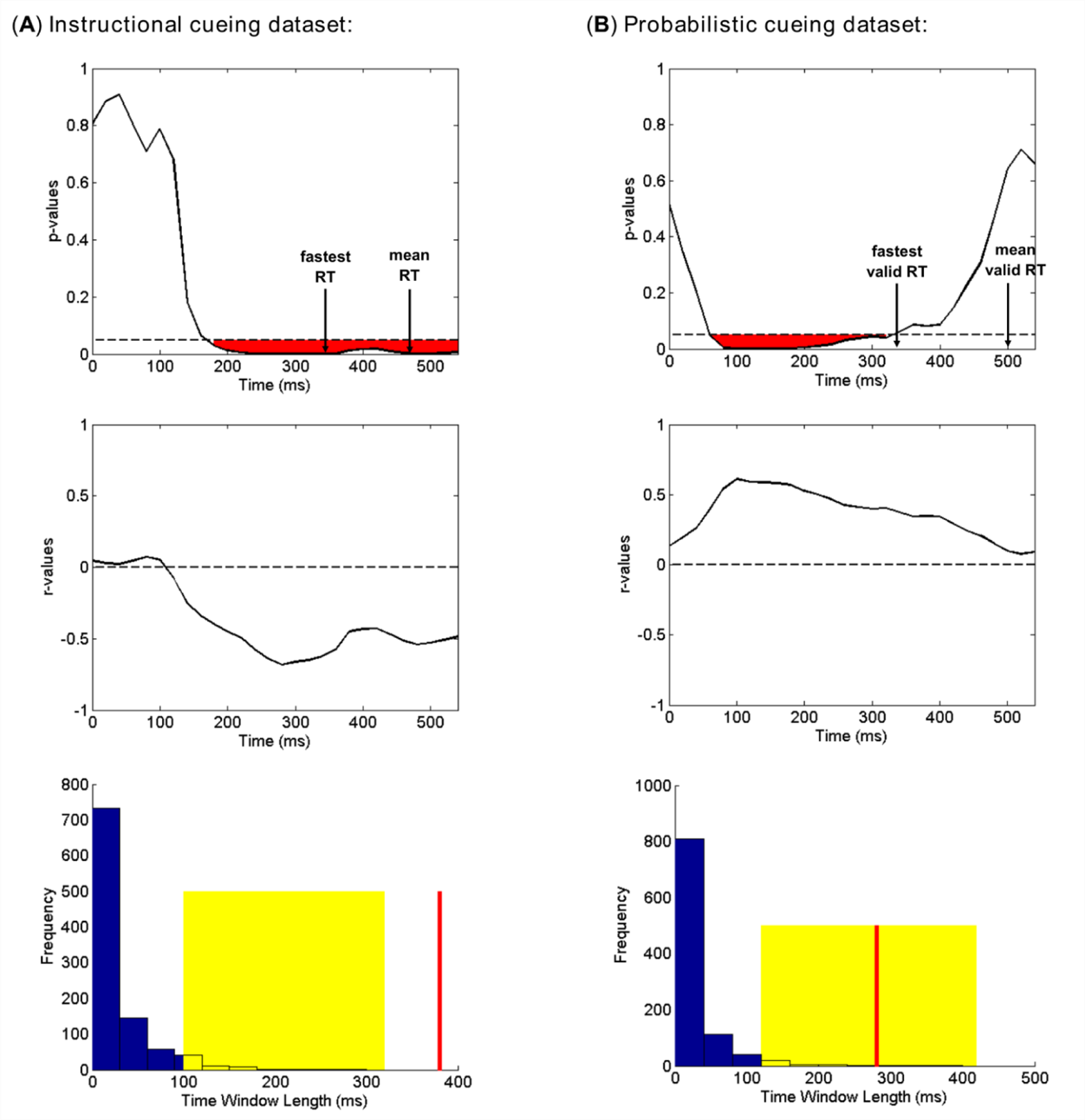
Results of between-subject correlation between target-related decoding accuracy and RT effects (*A*: RTs to attended targets; *B*: invalid RTs – valid RTs) in Instructional cueing dataset (*A*) and Probabilistic cueing dataset (*B*). For each dataset, the null distribution was estimated from 1000 permutations of the data, by randomly pairing one subject’s decoding accuracy with another subject’s RT effects. If the window length from the observed data (red line) falls within the top 5% of values from the null distribution (indicated by the yellow area), the observed window is considered to be significant.

Next, we correlated the univariate ERP difference waves (cued target *minus* uncued target, with left and right targets combined; see Figure S2 in Supplemental Information) with RT or RT difference across subjects in each dataset. The correlation analysis was performed at each time point within the 0-560 ms post-target interval (Fig. 9). FDR correction was first applied to correct for multiple comparison across channels, and permutation test was then applied to find significant clusters of contiguous time points. No significant cluster was identified for either dataset.

**Figure 9.**
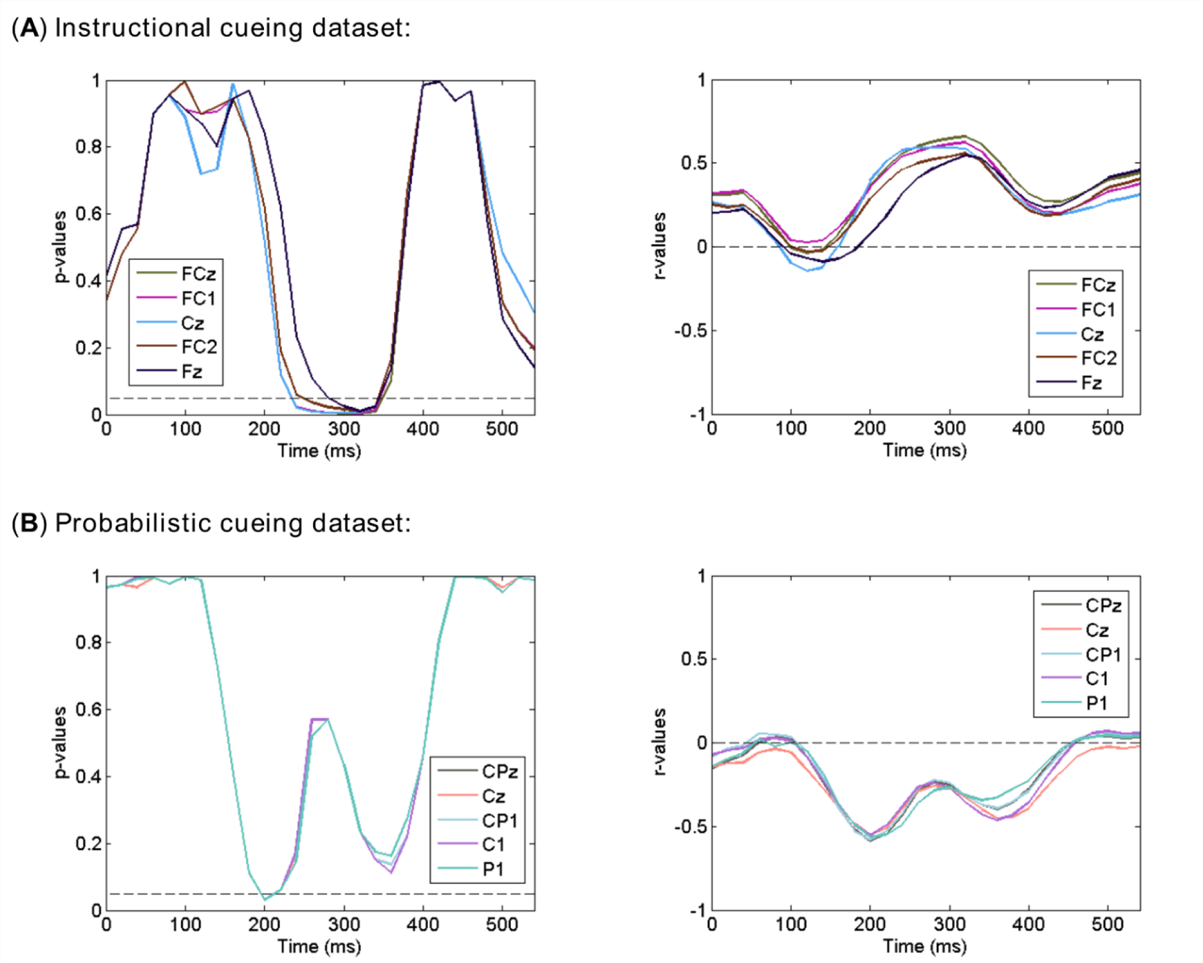
Results of between-subject correlation between target-related ERP difference wave (cued target *minus* uncued target) and RT effects (Panel *A*: RTs to attended targets; Panel *B*: invalid RTs – valid RTs) in Instructional cueing dataset (*A*) and Probabilistic cueing dataset (*B*). 5 channels with the smallest averaged *p* values within 0-560 ms interval were shown. All *p* values were FDR-corrected for channels at each time point. No significant effects were found.

## 4. Discussion

We applied machine learning approaches to multichannel ERP data to examine the dynamics and functional significance of neural representations of attention control and selection in two cued visual spatial attention experiments (probabilistic cueing vs. instructional cueing). SVM-based multivariate decoding was performed at each time point in the cue-related time period (cue left vs. cue right) and in the target-related time period (cued target vs. uncued target). We found that following cue onset, the decoding accuracy began to rise above chance level at ∼80 ms for the Instructional cueing dataset, and at ∼160 ms for the Probabilistic cueing dataset. Across subjects, decoding accuracy between ∼460 and ∼660 ms post-cue predicted the magnitude of attentional modulation of subsequent target processing indexed by the target-evoked N1 component, indicating that the attentional set or template implemented between ∼460 and ∼660 ms directly affected the attention selection of the target. During target processing, the decoding accuracy began to rise above chance level at ∼100 ms for the Instructional cueing dataset, and at ∼160 ms for the Probabilistic cueing dataset. Across subjects, decoding accuracy over a broad post-target time window predicted RT (or RT cueing effect), suggesting that the distinctness of neural representations of attended information affected subsequent behavioral performance. In contrast, univariate ERP analysis failed to provide an association between ERP attention effects during the cue-target interval and the attentional modulation of target-evoked response or between ERP attention effects during target processing and subsequent behavioral performance. Together, these findings suggest that multivariate decoding analysis of ERPs is a powerful approach, and along with the conventional univariate ERP analysis, offers more comprehensive insights into the neural mechanisms of attention control and selection in visual spatial attention.

### 4.1 Decoding attention control in the anticipatory attention

Consistent with our hypothesis that instructional cueing is associated with earlier formation of the attentional set, the decoding accuracy began to rise above chance level at ∼80 ms following the onset of instructional cue, but at a much delayed time of ∼160 ms following the onset of probabilistic cue. EDAN, an early ERP component thought to mark the initial attention shift toward the attended location (Harter et al., 1989; Hopf & Mangun, 2000), often appears at ∼200 ms post-cue [sometimes earlier at ∼160 ms, see Nobre et al. (2000)] in previous ERP analysis of the spatial cueing paradigms. The timing difference between decoding onset (∼80 or ∼160 ms) and EDAN latency (∼200 ms) suggested that the attention shift was initiated earlier than previously thought. One might argue that the different physical properties of visual stimuli used as cues (left vs. right) might explain the early decodability of the data. This is unlikely to be the case, because if this were the case, the decoding accuracy should rise above chance level equally early for both instructional cueing and probabilistic cueing, since similar cues were used in both paradigms. Although no study has explicitly compared the onset latency of EDAN between instructional cueing and probabilistic cueing, the ∼80 ms versus ∼160 ms timing difference in the onset of decodability between instructional cueing and probabilistic cueing, along with the higher decoding accuracy in instructional cueing than in probabilistic cueing, while not unexpected, was demonstrated here for the first time. This suggests that unilateral attention orienting (instructional cueing), compared with attention spread across both visual hemifields (probabilistic cueing), facilitate the early formation and enhance the distinctness of attentional set.

As time progressed, the decoding accuracy continued to rise and reached a local maximum around ∼300 ms in both datasets, and then declined slightly, but remained well above chance level until the end of the cue-target interval. This temporal pattern suggests that the subject, following instructions, was able to maintain a state of covert attention until the onset of target processing. In contrast to the relatively stable multivariate decoding dynamics, univariate ERP analysis showed that ADAN, an ERP component reflecting supramodal mechanisms of attentional engagement in frontal areas (Eimer et al., 2002), appeared at ∼350 ms in both datasets, whereas LDAP, an ERP component indexing increase in the excitability of occipital cortical neurons (Hopf & Mangun, 2000; Kelly et al., 2009), started to appear at ∼400 ms in Instructional cueing dataset and at ∼450 ms in Probabilistic cueing dataset (see Figure S1 in Supplemental Information). Both ADAN and LDAP vanished after ∼700 ms, and a contralateral pre-target negativity with a frontal concentration became the dominant ERP phenomenon. Although previous studies have already reported contralateral negativity throughout the pre-target period, the scalp distribution of this negativity varied across studies, i.e., within the occipital-parietal area (i.e., BRN) (Grent-’t-Jong et al., 2011; Grent-’t-Jong & Woldorff, 2007), the frontal area (Hopf & Mangun, 2000) or a broader area including frontal and parietal regions (Dale et al., 2008). This indicated that multiple neural processes might underlie the late negativity, making it insufficient to examine these processes in a univariate ERP approach. Despite this uncertainty, our multivariate decoding analysis suggested that the distinctness of neural representation of the attentional set during this late cue-target interval (> 700 ms) did not significantly decline compared to the early cue-target interval (< 700 ms).

Decoding accuracy is an indicator of how well attended information is represented in the brain. As such, it is reasonable to expect that better representation of attended information in the cue-target interval will result in stronger attentional modulation of target processing. We tested this hypothesis by correlating, across subjects, the decoding accuracy derived from multivariate classification analysis with target-evoked N1 attention modulation. As shown in Fig. 5, during the time period of 460-660 ms, decoding accuracy is positively correlated with the size of attention effects on the target-evoked N1 component. In visual spatial attention, this time period was often regarded as the critical stage in the implementation of the attention control state for the representations of task-relevant locations (Dale et al., 2008; Grent-’t-Jong & Woldorff, 2007; Hopf & Mangun, 2000). Our results thus suggest that the anticipatory attentional state that was implemented at this time can directly impact subsequent attention selection of behaviorally relevant stimuli, and more importantly, this attentional state was indexed as a whole brain ERP pattern instead of univariate ERP amplitudes based on any single electrode, as our univariate ERP analysis did not reveal any correlation between ERP amplitudes and target-evoked N1 attention modulation. The weight maps of the classifier during 400-600 ms interval showed a frontal-posterior pattern that corresponded with a combined ADAN and LDAP topography (see Fig. 4), further supporting the above notion. During the later part of the cue-target interval (>700 ms), although the decoding accuracy did not significantly decline, it no longer predicted N1 attention modulation. This contrasts with previous reports that late negativity within the occipital-parietal area (Grent-’t-Jong et al., 2011) or the frontal area (Dale et al., 2008) alone predicted N1 attention modulation. On one hand, the relatively easy discrimination task used in our experiments, compared with a more difficult task, could substantially decrease the amplitudes of the late negativity (Grent-’t-Jong et al., 2011), which might then reduce the decodability during the late cue-target interval. On the other hand, in the late cue-target interval, the neural representation involving multiple neural processes captured by the multivariate decoding analysis may not have the same simple relationship with N1 attention modulation. The above two aspects may underlie the reason that above chance level decoding in the late cue-target interval did not predict N1 modulation by attention.

### 4.2 Decoding attention selection during target processing

Following the onset of the target stimulus, the decoding accuracy began to rise above chance level at ∼100 ms and ∼160 ms for the Instructional and Probabilistic cueing datasets, respectively. This again illustrated that attention exerted an earlier influence on target processing under instructional cueing than under probabilistic cueing. Although no study has explicitly compared the onset of attention effects between instructional cueing and probabilistic cueing using univariate ERP analysis, previous studies have shown that attention selection of sensory processing occurred as early as P1 component at ∼100 ms after stimulus onset (Hillyard & Anllo-Vento, 1998; S. J. Luck et al., 2000). In this study, significant attention effect for P1 component was not observed in the univariate ERP analysis, but the multivariate decoding analysis shows that attention effects are represented in multivariate patterns during the same time period. Moreover, the higher decoding accuracy in instructional cueing relative to probabilistic cueing was again as expected, suggesting that unilateral attentional focus facilitates the attention selection of the target stimulus. As time progressed, further differences between the two paradigms emerged. In Instructional cueing dataset, the decoding accuracy remained relatively high (>0.8) after reaching the peak at ∼300 ms (Fig. 7A), while in Probabilistic cueing dataset, the decoding accuracy gradually declined after reaching the peak at ∼300 ms, and became non-significant near the end of the analysis period (Fig. 7B). This pattern was also expected from the paradigm requirements. In the probabilistic cueing experiment, subjects needed to orient their attention from the cued location to the uncued location upon seeing targets in the uncued location, which caused a gradual decrease of the decodability between valid and invalid targets, whereas in the instructional cueing experiment, no such attention shift was needed to complete the task, and the subject’s attention focused on the cued location even after the target stimulus appeared in the uncued location (such target stimuli were ignored). Utilizing the individual differences in decoding accuracy and behavioral performance, the functional significance of decoding accuracy was examined by correlating decoding accuracy during target processing and RT (or RT cueing effect) across subjects. Significant correlation was observed soon after target onset (see Fig. 8), and this correlation remained significant until the response was made, leading support to the notion that stronger attention selection of the target stimulus leads to better behavioral performance.

It is worth noting that from decoding analysis it is not easy to discern the individual contribution of distributed cognitive processes activated by target onset. Univariate ERP analysis has revealed attentional modulation of neural activity at multiple stages of information processing, including ERP differences following early sensory components (e.g., P1 and N1) as well as prior response execution (e.g., Nd1, Nd2 and LPD) (Curran et al., 2001; Eimer, 1996; Eimer, 1998; Mangun & Buck, 1998; Mangun & Hillyard, 1991). Although the precise functional correlates of these late ERP activities are still not clear, they should reflect perceptual, cognitive and motor consequences of spatial attention (Mangun & Buck, 1998). More interestingly, the maximum decoding accuracy at ∼300 ms roughly corresponds with the latency of Nd2 which is part of the broader LPD (Curran et al., 2001; Eimer, 1996; Eimer, 1998), indicating that the decoding accuracy might be primarily driven by Nd2. This inference was further supported by the similarity between the SVM weight maps (see Fig. 7) and ERP topographical maps (see Figure S2 in Supplemental Information). Despite these important findings, univariate ERP differences between cued and uncued targets at single electrode did not predict behavioral performance (see Fig. 9). One reason may be that the ERP patterns over the scalp could differ between different stages of information processing in spatial attention. By contrast, the decoding analysis can capture the ERP pattern over the whole scalp, which appeared to be a reliable predictor for behavioral performance in spatial attention tasks.

### 4.3 Conclusions

In the present study, we applied a machine learning approach to analyze neural representations of attention from multichannel ERP patterns over the whole scalp in two independent visual spatial attention experiments. Across the two experiments, the direction of covert attention can be decoded, and the decoding accuracy during the cue-target interval (∼460-660 ms post-cue) in which anticipatory attention set was implemented predicted the attentional modulation of target-related N1 amplitude. Also across the two experiments, after target appearance, attended targets can be decoded from unattended (or less attended) targets, and the decoding accuracy predicted behavioral performance (RT or RT cueing effect). However, no brain-behavior association was observed when the correlation analysis was based on univariate ERP analysis, i.e., ERP amplitudes from single channels. Therefore, our findings suggest that top-down attentional control and its modulation on target processing is more comprehensively represented in multichannel ERP patterns over the whole scalp, rather than ERP amplitudes measured in specific channels, and that naturally occurring individual differences in neural and behavioral variables enable the study of the functional significance of decoding accuracy derived from multivariate classification in attention control and selection.

## Acknowledgments

This work was supported by the National Science Foundation of China (No. 61601294, 61571295), NIH Grant R01 MH117991, Natural Science Foundation of Shanghai (No. 18ZR1432700), Research Foundation of Shanghai Municipal Health Bureau (No. 20174Y0020), Shanghai Jiao Tong University (No. YG2017MS44), Shanghai Mental Health Center (2016-QH-01, 2016-YJ-11) and Science and Technology Commission of the Shanghai Municipality (No. 13dz2260500). X. H. also gratefully acknowledges the support of Shanghai Jiao Tong University K. C. Wong Medical Fellowship Fund. The funding sources had no role in study design, data collection and analysis, decision to publish, or preparation of the manuscript.

