## Supplemental Figures for "Decoding Attention Control and Selection in Visual Spatial Attention"

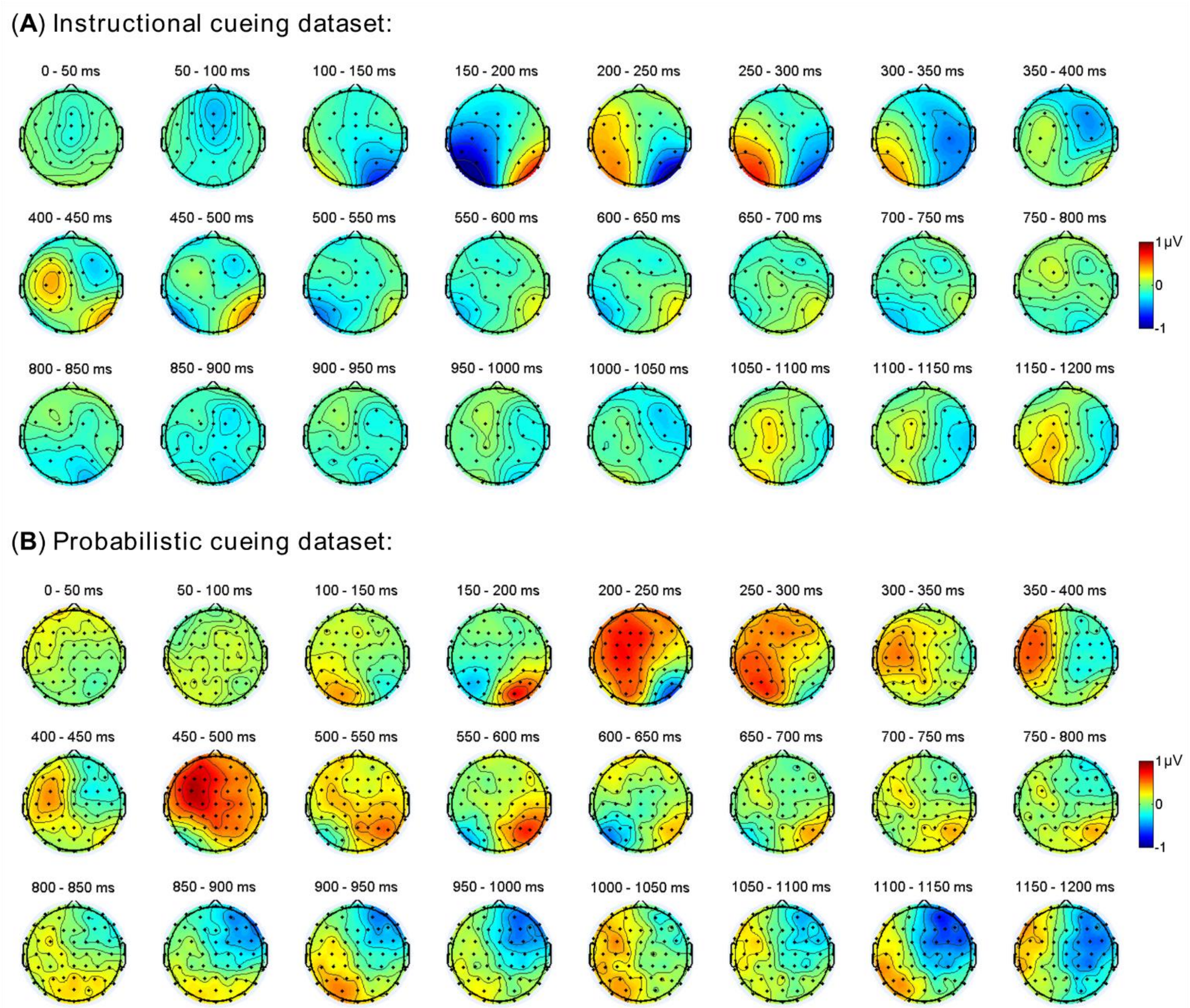


**Figure S1.** Topographical maps of ERP difference waves (cue left *minus* cue right) from successive time points within the indicated windows were averaged and shown for each dataset.

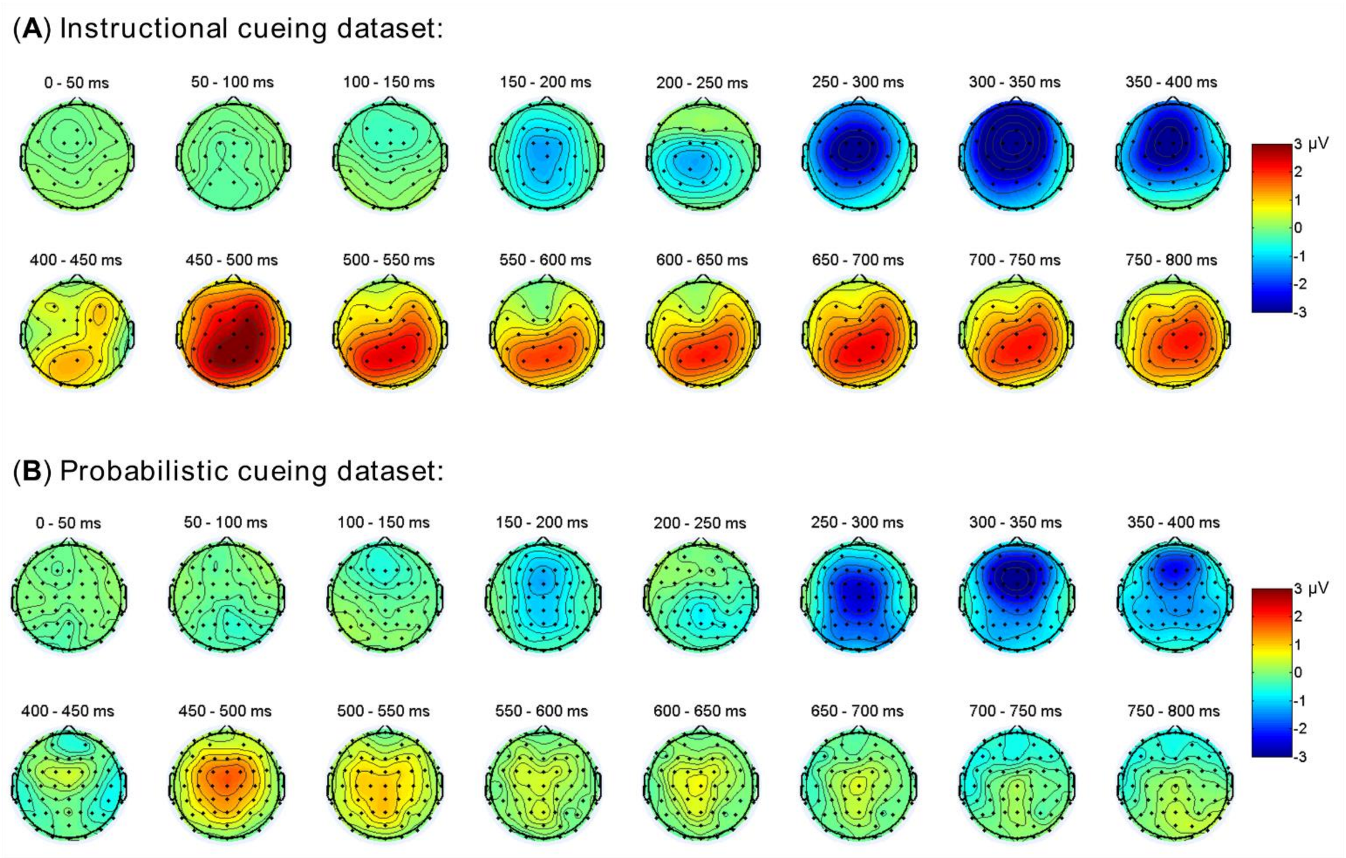


**Figure S2.** Topographical maps of ERP difference waves (cued target *minus* uncued target, with left and right targets combined) from successive time points within the indicated windows were averaged and shown for each dataset.
